# The molecular framework underlying the growth and defense tradeoff in *Arabidopsis* roots in response to danger signals

**DOI:** 10.1101/2024.04.26.591300

**Authors:** Souvik Dhar, Soo Youn Kim, Heeji Shin, Jongsung Park, Ji-Young Lee

**Author notes:** Author for correspondence: Ji-Young Lee. **One sentence summary:** STZ and its relatives finetune the root growth and stress resistance in response to an endogenous danger signal. The author **responsible** for distribution of materials integral to the findings presented in this article in accordance with the policy described in the Instructions for Authors (https://academic.oup.com/plcell/) is: Ji-Young Lee.

## Abstract

Elevated stress signaling often compromises plant growth by suppressing proliferative and formative divisions at the meristem. Plant Elicitor Peptide 1 (PEP1), an endogenous danger signal triggered by both biotic and abiotic stresses in *Arabidopsis thaliana*, suppresses proliferative divisions, alters xylem vessel organizations, and disrupts cell-to-cell symplastic connections in the root. To gain an insight into the dynamic molecular framework that modulates root development under elevated danger signals, we performed a time course RNA-sequencing analysis at the root meristem following PEP1 treatment. A series of data analyses revealed that STZ and its homologs are a potential nexus between the stress response and proliferative cell cycle regulation. We observed that STZ differentially controls the cell cycle, cell differentiation, and stress response genes at various tissue layers in the root meristem through various functional, phenotypic, and transcriptomic analyses. Our study further indicated that the expression level of *STZ* in response to stresses is critically important to enable the growth and defense tradeoff. The findings here provide valuable information about the dynamic gene expression changes upon perceiving danger signals at the root meristem and future engineering schemes to generate stress-resilient plants.

## Introduction

As sessile organisms, plants use distinctive strategies from animals when they encounter biotic and/or abiotic stresses. Plants quickly turn to investment in defense/stress tolerance programs for survival at the expense of growth (Huot et al., 2014). For this transition, stress tolerance, starting with a complex exchange of plant-environmental signals at a cellular level, should propagate to the entire organ and body and suppress the growth program in meristems (Figueroa-Macias et al., 2021). Understanding the dynamics of perception and propagation of stress signals and their downstream transcriptional framework would help find how plants balance growth and stress responses and develop sustainable crops under environmental changes.

Biotic and abiotic stress signals from the outside trigger the production of so-called ‘danger signals’ in the cell. Danger signals in the form of small metabolites or peptides are secreted outside the cell and sensed by neighbors, thereby further turning on stress responses. Plant elicitor peptides (PEPs) are one such danger signals that are activated by both biotic and abiotic stresses (Huffaker et al., 2006; Huffaker et al., 2013; Ross et al., 2014; Yamada et al., 2016; Nakaminami et al., 2018; Hander et al., 2019). In *Arabidopsis thaliana* (hereafter Arabidopsis), synthetic PEP1 treated to seedlings was shown to suppress apical root growth, trigger the formation of excessive root hairs, and modify vascular tissue arrangement within the stele (Poncini et al., 2017; Dhar et al., 2021; Okada et al., 2021). However, how PEP1 transcriptionally modulates root development and whether this is an indispensable part of the tradeoff between growth/development and stress resistance/defense, which we collectively call as ‘the growth and defense tradeoff’ are yet to be elucidated.

Transcriptional gene regulatory networks (GRNs) are essential in stress responses at various levels. Many transcription factors (TFs) in stress responses have been identified, and GRNs involving these TFs are under active investigation (Han et al., 2020b). Among more than 2000 TFs in Arabidopsis, the zinc finger proteins form one of the largest TF families and are divided into various sub-families based on the position of the Cys and His residues in their secondary structures (Miller et al., 1985; Klug, 2010; Han et al., 2014). The C2H2-type zinc finger TF family, which includes 176 genes in Arabidopsis (Englbrecht et al., 2004; Xie et al., 2019), has been reported to function in plant growth and development, biotic and abiotic stress resistance (Han et al., 2020a) and programmed cell death (PCD) (Feng et al., 2023).

In this study, we employed PEP1 as a mediator triggering a broad range of biotic and abiotic stresses and then investigated GRNs downstream of PEP1 to find key TFs in controlling stress responses and root growth and development. Our investigation suggested that SALT TOLERANCE ZINC FINGER (STZ/ZAT10) and related C2H2 zinc finger TFs including ZAT6 and AZF3 might serve as nexus between stress response and root apical growth. PEP1 rapidly induces the expression of genes encoding these TFs in the root apical meristem, including a stem cell niche. Transgenic roots overexpressing *STZ* phenocopied the roots treated with PEP1, displaying the suppression of root apical growth and ectopic vascular tissue differentiation (Dhar et al., 2021). This phenotype was consistent with changes in transcriptomes in response to STZ in the root meristem, in which cell cycle genes were suppressed while developmental and environmental programmed cell death (PCD) genes as well as stress response TFs were activated. Suppression of cell cycle genes required more STZ dosage than the activation of stress response TF genes. These findings have the potential to guide sustainable crop plant growth under stress conditions, which might be possible by fine-tuning the expression of *STZ* and its homologs.

## Results

### PEP1-mediated transcriptional changes in the root meristem

PEP1 drastically suppresses the apical root growth by inhibiting proliferative cell divisions (Poncini et al., 2017; Jing et al., 2019; Dhar et al., 2021). We observed a clear reduction of the meristem size as early as 6h after PEP1 treatment (Figure S1a and b). We also reported that PEP1 influences formative divisions of the vascular initials in the meristem (Poncini et al., 2017; Jing et al., 2019; Dhar et al., 2021). To understand the genes and regulatory pathways PEP1 directs for cellular reprogramming in the root meristem, we performed a time-course transcriptome profiling for PEP1-treated root meristems (Figure 1a). Root meristems were dissected from 5-day-old Arabidopsis Col-0 seedlings that were treated with 1μM of synthetic PEP1 for 3, 6, 12, and 24 hours, respectively. Total RNAs extracted from these samples were processed for sequencing using Illumina NovaSeq6000 system (Figure S1).

**Figure 1.**
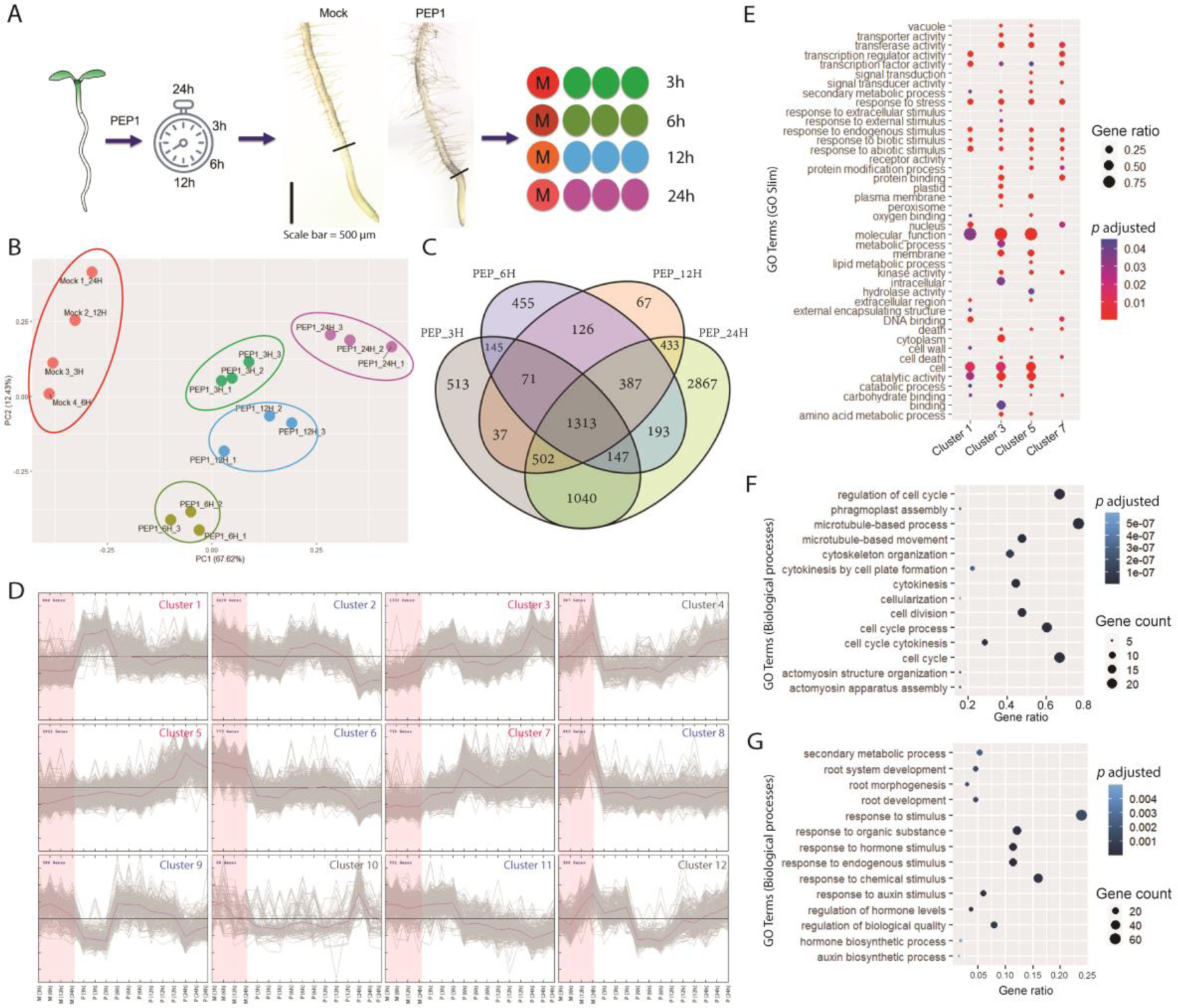
Identification of PEP1-influenced meristem-specific regulatory cascades through a time-course RNA sequencing. **(A)** A schematic plan of the RNA-seq experiment. We used four different time points (3h, 6h, 12h, and 24h) to harvest meristem samples used for RNA isolation. **(B)** A principal component analysis (PCA) of 16 RNA-seq samples based on the top 5000 most variable genes. **(C)** A comparison of differentially expressed genes [DEGs; false discovery rate (FDR) < 0.01, |fold change| ≥ 1.5) identified in this work. The comparisons were made between 3h, 6h, 12h, and 24h post-PEP1 treated root meristem samples. In total, 8,296 unique DEGs identified are presented in Table S3. **(D)** K-means clustering of 8,296 DEGs were grouped into 12 clusters (also presented in Table S4). M denotes mock and P denotes PEP1 treatment. **(E)** Gene ontology (GO Slim) enrichment analysis (FDR < 0.05) of the PEP1-induced DEGs from clusters 1, 3, 5, and 7 from panel D. The total number of DEGs among these clusters and the respective GO term (biological processes) enrichment analysis are presented in Table S6. **(F** and **G)** The dot plots represent gene ontology (biological processes) enrichment analysis (FDR < 0.01) of the PEP1 suppressed DEGs from clusters 9 and 11 from panel D. Notably, among the PEP1 suppressed clusters, 9 and 11 were enriched with cell cycle and root development related DEGs, respectively. Top 14 GO terms of each cluster were displayed. Detailed analysis for each clusters are in Table S5.

Over 800 million high-quality clean RNA-seq paired-end reads were generated from 16 samples (Table S1; methods). The expression levels (RPKM) of all annotated transcripts are provided in Table S2. A principal component analysis (PCA) of the top 5000 most variable genes revealed that the mock samples’ integrity remained distinctive enough to be separated from the PEP1 treated samples by PC1 even though each of mock samples was collected at different time points (Figure 1b). We identified a total of 8296 differentially expressed genes (DEGs; fold change [FC] ≥ 1.5 and false discovery rate [FDR] < 0.01) in PEP1 treated samples compared with the mock samples (Figure 1c; Table S3). Unique DEGs significantly increased 24h after PEP1 treatment (Figure S2; Table S3). To detect PEP1-dependent transcriptome reprogramming over time, we clustered expression patterns of 8296 DEGs following K-means algorithms (Lloyd, 1982) (Figure 1d; Table S4). Out of 12 clusters, clusters 1, 3, 5, and 7 exhibited PEP1-induced “rapid transient”, “rapid stable”, “late”, and “late stable” expression dynamics, respectively. Among the PEP1-repressed clusters, cluster 2 showed “late”, clusters 6 and 8 showed “rapid stable”, cluster 9 exhibited “rapid transient”, and cluster 11 displayed “late stable” suppression.

To learn biological processes affected by PEP1 in the root meristem, we conducted a gene ontology (GO) term enrichment analysis (FDR ≤ 0.01) for the above 9 clusters, showing explicit expression dynamics in response to PEP1 (Table S5). Because of the high volume of over-represented GO terms, we performed GOSlim categorization for the PEP1-induced clusters to gain a broader overview of GO contents (Figure 1e, Table S6). As expected, clusters with genes induced by PEP1 were enriched with genes involved in responses to various biotic and abiotic stimuli. Among the five clusters with genes downregulated under PEP1 treatment, clusters 9 and 11 were enriched with genes that regulate cell cycle and root development, respectively (Figure 1f and g). The Cluster 9, composed of 388 DEGs, displayed a significant enrichment of “regulation of cell cycle (GO:0051726)”, “cell cycle (GO:0007049)”, and “cell division (GO:0051301)” related GO terms (Figure 1f, Table S5). Moreover, the GO term analysis results from cluster 11 revealed the presence of significantly enriched DEGs involved in root development (GO:0048364) and morphogenesis (GO:0010015) (Figure 1g, Table S5). Since root meristem and growth are perturbed in exposure to PEP1 (Figure S1; Dhar et al., 2021), we focused on clusters 9 and 11 among the PEP1-repressed clusters in further study.

Next, we visualized the root cell type-specific expression patterns for genes belonging to clusters 9 and 11 (Brady et al., 2007; Zhang et al., 2019). Most genes in cluster 9 that PEP1 rapidly suppressed showed enriched expressions deep inside the root, such as xylem (S4 and S18) (Figure S3a). Genes in cluster 11, enriched with genes in root development and morphogenesis, showed differential expression in varying cell types (Figure S3b).

To identify the TFs induced by PEP1, we combined all DEGs from either “transcription regulator activity (GO:0030528)” or “transcription factor activity (GO:0003700)” terms, in clusters 1, 3, 5, and 7 (Figure 1e, S4a and b; Table S7). 344 TFs were found and classified into 43 distinct classes based on the protein domains using the InterPro database (https://www.ebi.ac.uk/interpro) (Figure S4c; Table S8). We observed that the stress-induced ERF, MYB, WRKY, and Zinc finger (ZF) domain TFs are enriched in the PEP1-induced clusters. Their root cell-type expression patterns (Brady et al., 2007; Zhang et al., 2019) indicated significant enrichment of the PEP1-induced transcriptional regulations inside the stele (Figure S4d).

### Identification of candidate transcription factors controlling the PEP1-dependent reprogramming of root development

To identify key transcription factors that trigger the reprogramming of root development in response to PEP1, we further investigated 344 TFs induced by PEP1 (Figure S4, Table S7). Employing STRING database version 12.0 (https://string-db.org/; (Szklarczyk et al., 2015)), we queried the interaction networks among these 344 TFs (interaction score > 0.7). 225 interactions among 159 TFs were identified (Table S9). In the most prominent network composed of 94 TFs with 162 interactions, *SALT TOLERANCE ZINC FINGER* (*STZ*), *COLD INDUCED ZINC FINGER PROTEIN 2* (*ZAT6*), *RESPONSIVE TO HIGH LIGHT 41* (*ZAT12*), *WRKY33*, *WRKY40*, and *DERB2A* were found as the most highly connected TFs (Figure S5a).

PEP1 rapidly suppresses the apical root growth likely by repressing cell cycle progression (Figure S1). This is indicated by the overrepresentation of cell cycle-related genes in cluster 9 (Figure 1f). Consistently, our assessment of cell proliferation using a PlaCCI, Plant Cell Cycle Indicator, (Desvoyes et al., 2020) line indicated a significant reduction of the markers related to the cell division (CYCB1;1 and CDT1a) in the meristem treated with PEP1 for 24h (Figure S5b). Since PEP1-mediated suppression of cell proliferation was clearly observed within the time scale of our transcriptome data, we asked whether PEP1-induced TFs suppress cell cycle regulators. To address this, we searched for the connectivity between 344 PEP1-induced TFs with 388 DEGs from the cell cycle cluster 9 in the STRING database. We kept the interaction parameter as “high confidence (0.700)” to avoid the large volume of noise. This *in silico* analysis revealed a large network composed of two densely connected subnetworks (Figure 2a; Table S10): one with a PEP1-induced TF cluster (Figure S5a) and the other with a distinct cell cycle gene cluster (Figure 2a). Interestingly, these two subnetworks were connected via DOF3.7 (At3g61850), PIF1 (At2g20180), and ABI5 (At2g36270), which then connected to WRKY40, DREB2A, WRKY33, STZ and ZAT12, which serve as the core TFs in the PEP1 induced TF cluster.

**Figure 2.**
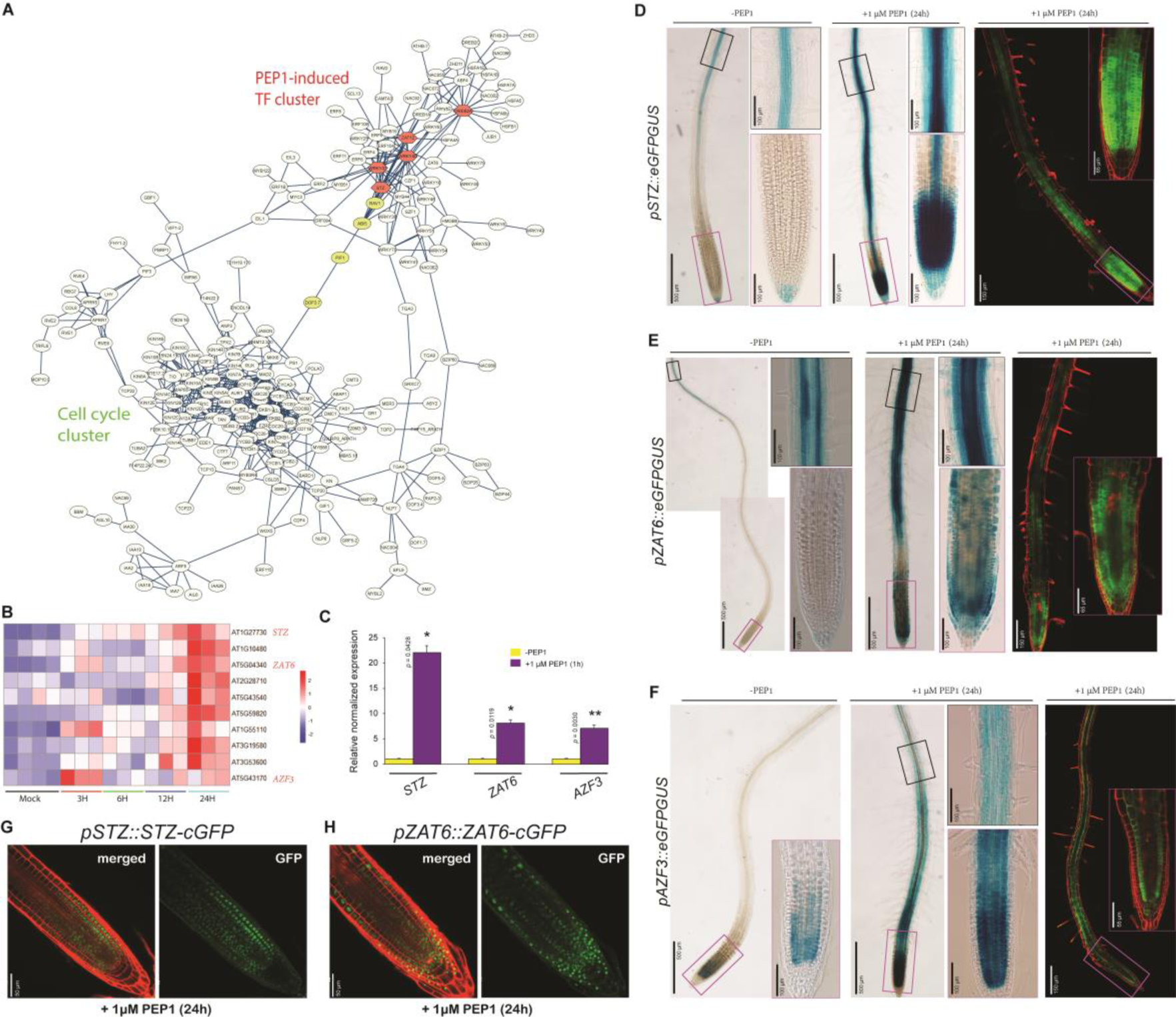
STZ and related C2H2 zinc finger TFs are a potential nexus between PEP1-induced and suppressed genes. **(A)** An interaction network of the PEP1-induced TFs together with the PEP1-suppressed cell cycle cluster as inferred from the STRING database version 12.0 (interaction score > 0.7) (Szklarczyk et al., 2015). The interaction scores are in Table S10. The network was generated at Cytoscape (www.cytoscape.org) (version: 3.10.1) using the above interaction score. **(B)** An expression heatmap showing the transcription patterns of the 10 C2H2 zinc finger (ZF) TFs presented in Table S8. These ZF TFs showed an early induction after the seedlings perceived PEP1. **(C)** The expression levels of three selected TFs as analyzed by qRT-PCR. The seedlings were treated with 1 M PEP1 for 1 hour (1h), and the roots were dissected to extract total RNAs for gene expression analyses. For each sample, at least three technical replicates were run, and the fold changes were determined compared to the control sample (-PEP1). Expression values were presented as ± SEM. Statistical significance of differences was determined by Student’s *t*-test (\*\**P* ≤ 0.01, \**P* ≤ 0.05). **(D - F)** Expression patterns of *STZ*, *ZAT6*, and *AZF3* in the 5 DAT wild type Col-0 plant without or with 1 M PEP1 treatment for 24 hours. The transcriptional expression patterns of these genes were visualized with the GUS or GFP system (*pSTZ::eGFPGUS*, *pZAT6::eGFPGUS*, *pAZF3::eGFPGUS*). Notably, the expression of these three ZF TFs were enriched in the meristem at 24h post-PEP1 treatment. **(G** and **H)** The proteins of *STZ* (STZ-cGFP) and *ZAT6* (ZAT6-cGFP) were localized in the meristem as visualized after 1 M PEP1 treatment for 24 hours using the respective translational fusion lines (*pSTZ::STZ-cGFP* and *pZAT6::ZAT6-cGFP*). No GFP localization signal was observed in the meristem of the samples not treated with PEP1 (data not shown). Intriguingly, the ZAT6 localization is enriched at the stem cell niche (SCN).

In Arabidopsis, STZ, a C2H2 zinc figure (ZF) TF, has been reported to be involved in the multiple stress response pathways (Sakamoto et al., 2004; Mittler et al., 2006; Xie et al., 2012; Hoang et al., 2020; Feng et al., 2023). STZ was originally identified as a strong transcriptional repressor significantly influenced by abiotic stresses, especially salinity, cold, and draught (Lippuner et al., 1996; Sakamoto et al., 2000; Sakamoto et al., 2004). Among genes encoding C2H2 ZF TFs, *AZF3* (AT5G43170) and *ZAT6* (AT5G04340) are closely related to *STZ* (Figure S5c; (Xie et al., 2019)). In our transcriptome data, the expression of *AZF3*, *STZ*, and *ZAT6* was rapidly induced by the PEP1 exposure and then stayed at high levels (Figure 2b). Our quantitative real-time PCR (qRT-PCR) assay indicated the rapid induction of *STZ*, *AZF3*, and *ZAT6* transcription within 1h of PEP1 treatment (Figure 2c). Constitutive expression of *STZ* or *ZAT6* has been shown to confer the suppression of shoot and root growth in the Arabidopsis seedlings (Sakamoto et al., 2004; Shi et al., 2014), which is reminiscent of the phenotype of seedlings exposed to PEP1. Thus, we asked whether the *STZ* and its closely related C2H2 ZF TFs function as a nexus between PEP1 sensing and the reprogramming of the root development.

We next examined the tissue-specific expression patterns of *STZ*, *AZF3* and *ZAT6* in 5-day-old Col-0 seedling roots introduced with *pSTZ::eGFPGUS*, *pAZF3::eGFPGUS* and *pZAT6::eGFPGUS*, respectively (Figure 2d - f). The transcriptional reporter lines of *STZ* and *ZAT6* (*pSTZ::eGFPGUS* and *pZAT6::eGFPGUS*) did not show any expression of eGFP in the root meristem without PEP1 treatment, and GUS staining of *pSTZ::eGFPGUS* lines showed dim expression in the columella (Figure 2d and e). By contrast, in the seedling roots exposed to PEP1 for 24h, very high expression of *pSTZ::eGFPGUS* and *pZAT6::eGFPGUS* was observed throughout the meristem and in the stele of the elongation and differentiation zones (Figure 2d and e, S5e and f). Similar to *STZ* and *ZAT6*, we found an enhancement of *eGFP-GUS* expression in the meristem of the transcriptional reporter lines of *AZF3*, under PEP1 treatment (Figure 2f). The expression domain of the PEP1-induced *pZAT6::eGFPGUS* partly overlapped with PEP1-influenced *STZ* expression, whereas the expression domain of *pAZF3::eGFPGUS* in the meristem was mainly restricted to the cortex and endodermis (Figure 2e and f). Previous cell type-specific root map data (Brady et al., 2007; Zhang et al., 2019) predicted that these three ZF TFs are mainly expressed in the mature cambium and phloem companion cells in the root differentiation zone in unperturbed conditions (Figure S5d). Consistent with the prediction, we found the transcription of *STZ* and *ZAT6* within the stele of the differentiation zone, which was further induced by PEP1 (Figure S5e and f). By contrast, the transcription of *pAZF3::eGFPGUS* under the influence of PEP1 was mostly enriched in the endodermis and cortex layers (Figure S5g). Consistent with transcriptional reporter lines, we found the presence of STZ and ZAT6 proteins in the root meristem following the treatment with PEP1 (Figure 2g and h). The protein localization domains significantly overlapped between ZAT6 and STZ; however, ZAT6 seemed more enriched in the stem cell niche than STZ (Figure 2g and h). Collectively, these results implied that *STZ* and its close homologs that are ectopically induced in the meristem by PEP1 might coordinate the PEP1-mediated reprogramming of the root development.

### STZ reprograms root meristem as part of the PEP1 signaling cascade

PEPs are endogenous elicitors secreted in response to attacks by bacteria, fungi, and herbivores (Yamaguchi and Huffaker, 2011; Bartels and Boller, 2015). PEP1-6 are sensed by the LRR-RLK PEP1 RECEPTOR1 (PEPR1) and PEPR2. To confirm whether the STZ-mediated growth suppression network is downstream of the PEP1 signaling cascade, we first monitored the expression levels of *STZ* and *PEPR*s in the root at 1h post-PEP1 treatment in the *pepr1 2* and *stz* mutant (Figure S6). Consistent with the previous report (Jing et al., 2019), short-term treatment (∼ 1h) with 1 M PEP1 caused induction of ∼12 and ∼30-fold of *PEPR1* and *PEPR2* levels, respectively, in the WT seedling root (Figure S6a and b). In the *stz* mutant, the *PEPR1* and *PEPR2* were also highly induced by PEP1 (Figure S6a and b). However, in the *pepr1 2* double mutant, we could not observe *STZ* induction as high as in WT seedlings in response to 1hr PEP1 treatment (Figure S6c). These findings suggested that the PEP1-mediated signal perception through PEPRs is upstream of the STZ regulatory networks.

To reveal the role of *STZ* in the root growth and development in response to PEP1, we developed *STZ*-overexpressing transgenic lines using a well-established estrogen-inducible XVE system (Zuo et al., 2000; Siligato et al., 2016). We transformed Col-0 wild-type plants with the vector carrying *p35S::XVE>>pLexA::STZ* transgene. After selection by Basta resistance, the induction levels of *STZ* in the T2 generation of *STZ*-Overexpression (*STZ-OE*) plants were estimated by quantitative real-time PCR (qRT-PCR) at 24h post-estradiol treatment (Figure S7a). We selected three *STZ-OE* lines (lines 2, 6, and 9 with 3883.3-, 730.7-, and 263.8-fold *STZ* expression compared to Col-0) having very high to relatively low induction levels of *STZ* expression. When grown on a half-strength Murashige and Skoog (MS) medium, the T2 generation of *STZ-OE* lines #2, #6, and #9 displayed comparable root growth to Col-0. However, when grown in 10 M estradiol, seedlings of lines #2 and #6 displayed a significant reduction of root growth with pale yellow cotyledon phenotype (Figure S7b). Contrary to these, line #9 exhibited comparatively better root growth on estradiol media (Figure S7b). We further confirmed the root growth phenotype of these three independent transgenics in T3 generation (Figure 3a). These *STZ-OE* transgenic plants showed similar seedling and root growth phenotypes to those of the T2 generation (Figures 3b and c). Consistent with the severe root growth retardation, we observed an exhaustion of meristem cells in the *STZ-OE* line #2-5. This phenotype contrasts with the phenotype of line #9-6, which exhibited the presence of a pool of meristem cells after the long-term *STZ*-induction (Figure 3d and e, S8). In addition, when we turned off the *STZ* overexpression in 7-day-old seedlings by transferring them to a half-strength MS plate without estradiol, root growth in *STZ-OE* #9-6 line was fully restored, similar to Col-0 seedlings. However, we did not find root growth recovery in *STZ-OE* #2-5 and #6-3 (Figure S9). These results suggest that STZ can negatively regulate cell division and stem cell maintenance in the root meristem in a dosage-dependent manner.

**Figure 3.**
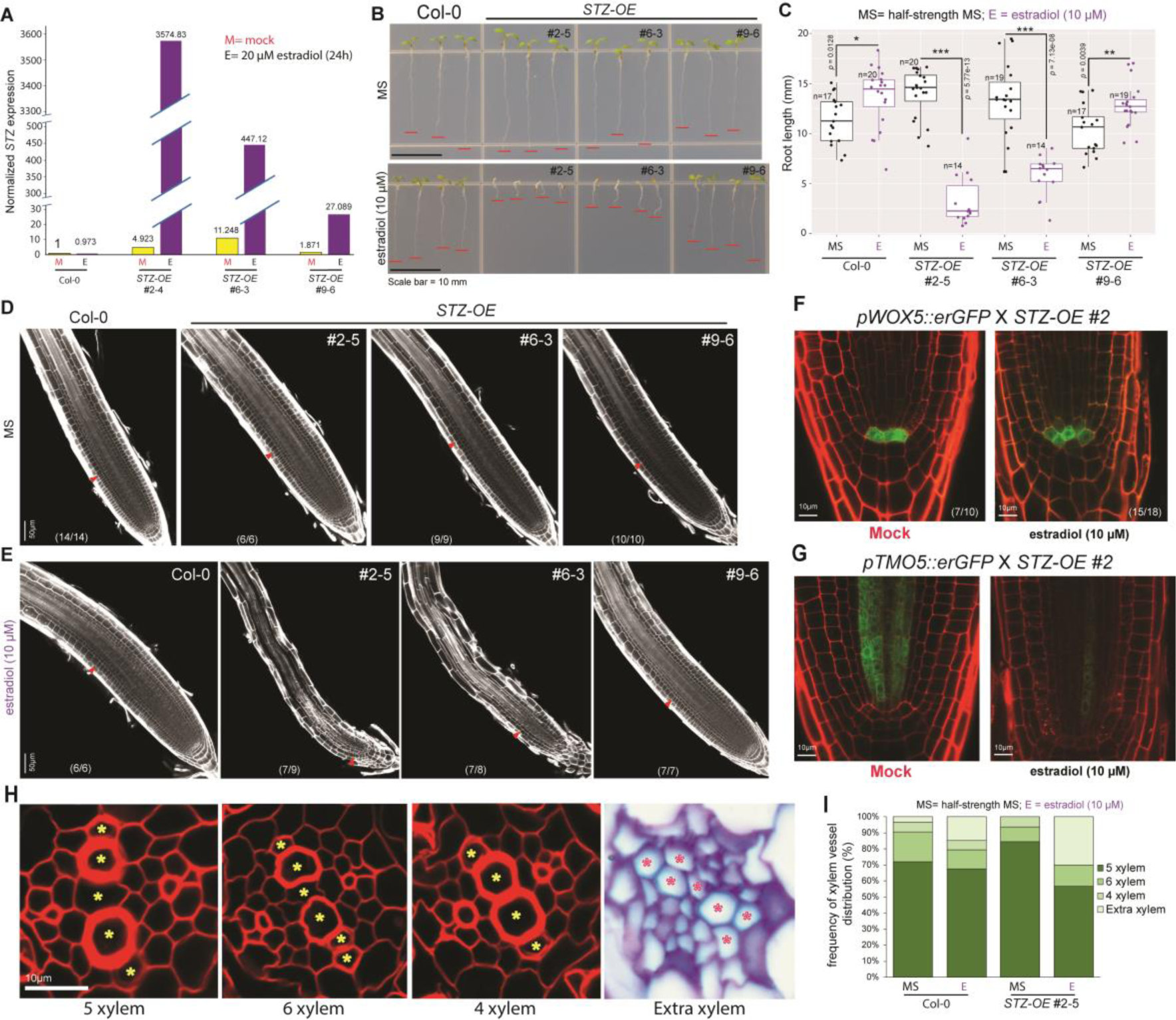
Overexpression of *STZ* regulates growth and differentiation in a dosage-dependent manner. **(A)** The expression level of *STZ* in *STZ-OE* lines after 24 hours of estradiol treatment as analyzed by qRT-PCR. The mock (ethanol) treated sample was used as a control to determine the fold change. The expression values were normalized by GAPDH and are presented as the ± SEM from three technical replicates. **(B)** The root growth of the 5 DAT old *STZ-OE* seedlings grown on either half-strength MS (MS) or half-strength MS media supplemented with 10 M estradiol. Red lines indicate the end part of the root tip. **(C)** The quantification of the root growth of *STZ-OE* seedlings grown on either half-strength MS (MS) or half-strength MS media supplemented with 10 M estradiol. The statistical significance of differences was determined compared to corresponding MS grown sample by Student’s *t*-test (\*\**P* ≤ 0.001, \*\**P* ≤ 0.01, \**P* ≤ 0.05). (**D** and **E**) The meristem growth of the Col-0 and three different *STZ-OE* lines. The MS-grown seedlings exhibited similar meristem growth between genotypes (**D**), whereas *STZ-OE* #2-5 and #6-3 roots displayed a significant reduction in the proliferative cell divisions in the meristem. The red arrow demarcates the junction between meristem and transition zone. The number in parenthesis of each panel indicates samples with similar results to the total independent root samples analyzed. (**F**) Identity of the quiescent center (QC) was deformed under the influence of *STZ* overexpression. The imaging was done using “stunted-root” growth phenotype plants from the progeny of F1 plants (F2 generation) crossed between *pWOX5::erGFP* and *STZ-OE*. Imaging was done at 5 DAT condition. Mock = seedlings grown on ½ MS media supplemented with ethanol; estradiol (10 M) = seedlings grown on ½ MS media supplemented with 10 M estradiol. The number in parenthesis of each panel indicates samples with similar results to the total independent root samples analyzed. **(G)** *STZ* overexpression significantly suppressed *pTMO5::erGFP* transcription in the 5 DAT old seedlings. **(H)** Representative images of four typical xylem arrangements in the Col-0 and *STZ-OE* #2-5 roots; the “5 xylem” cells phenotype, “6 xylem” cells phenotype, “4 xylem” cells phenotype, and “extra xylem” cells phenotype. The yellow and red asterisks indicate differentiated xylem vessels. **(I)** Quantification of the xylem differentiation phenotypes in the 5 DAT old Col-0 and *STZ-OE* #2-5 roots grown either on half-strength MS or half-strength MS supplemented with 10 M estradiol. The section images and xylem vessel distribution frequency calculation from different growth combination used are in Figure S11 and Table S11.

We then inquired whether the quiescent center (QC) and xylem precursors remained intact in the roots under a high dosage of *STZ*. By genetic crosses, we introduced transgenic constructs of *pWOX5::erGFP* (Sarkar et al., 2007), which marks the QC identity, and *pTMO5::erGFP* (Lee et al., 2006), the marker for xylem precursor cells, into the *STZ-OE* line #2. In the following F2 population, we monitored the expression patterns of those marker genes in 5-day-old seedlings grown on estradiol media (Figure 3f and g). *WOX5* domain frequently expanded beyond the original QC position (Figure 3f; S10), while the TMO5 expression was reduced drastically (Figure 3g). We also scored xylem differentiation patterns in the 5-day-old seedling roots of *STZ-OE* line #2-5 grown on 10 M estradiol media in comparison with WT (Figure 3h and i; S11) (Zhou et al., 2013; Dhar et al., 2021; Seo and Lee, 2021). In the media without estradiol, WT seedling roots showed “5 xylem cells” as the most prevalent type (∼72%), followed by the “6 xylem cells” (∼19%), “4 xylem cells” (∼6%) and the “extra xylem” (∼3%) types (Figure 3i; S11a; Table S11). The *STZ-OE* line #2 showed xylem patterning similar to the WT with the presence of 84% “5 xylem cells”, followed by 9% of “6 xylem cells” and 6% of “4 xylem cells” type (Figure 3i; S11b; Table S11). However, estradiol-induced *STZ-OE* line #2 displayed a significant increase of “extra xylem” vessel differentiation as compared to the WT seedlings grown on 10 M estradiol plates (∼15% in WT and ∼30% in *STZ-OE*#2) (Figure 3i; S11c and d; Table S11). These *STZ-OE* phenotypes recapitulate the PEP1-influenced phenotypes of WT, supporting the crucial role of STZ in the PEP-mediated reprogramming of root development (Dhar et al., 2021).

### Identification of downstream regulators of STZ acting in response to PEP1

To understand how *STZ* transcriptionally modulates the downstream signaling cascades, we generated RNA-seq data for the roots of *STZ-OE* #2-4 line sampled in 6h and 24h post-*STZ* induction (Figure S12a). The RPKM values of all mapped transcripts are provided for 9 samples with over 500 million high-quality paired-end reads (Table S12, S13). The PCA analysis with the top 5000 variable genes revealed the sample integrity between replicates (Figure 4a). At 1.5-fold cut-off we identified a total of 8,435 DEGs compared with the mock treated samples (Figure S12b and c; Table S14). At 24h post *STZ* induction, we observed more DEGs than 6h induction (Figure S12c). Although STZ acts as a transcriptional repressor, more than 50% of the DEGs were upregulated after STZ induction (Figure S12c). K-means clustering with these DEGs revealed 6 clusters (Figure 4b) with clusters 1, 5, and 6 showing suppression by STZ and clusters 2, 3, and 4 showing rapid to late induction by STZ (Figure 4b, Table S15). Since proliferative cell divisions in the root meristem were suppressed by both *STZ* overexpression and PEP1 treatment, we searched for the cell cycle genes by overlaying genes in a PEP1-suppressed cluster with genes suppressed by STZ (Cluster 1, 5 and 6 for *STZ-OE* data and Cluster 9 of PEP1 data; Table S4 and S15). 61 overlapping genes (Table S16, Figure S12d) belonged to the GO terms, including “regulation of cell cycle (GO:0051726)” and “regulation of mitotic cell cycle (GO:00007346)” (Figure 4c, Table S17). Interestingly, the expression of these 61 genes is significantly enriched in the developing xylem precursor (S4) cell type (Figure 4d; Table S18).

**Figure 4.**
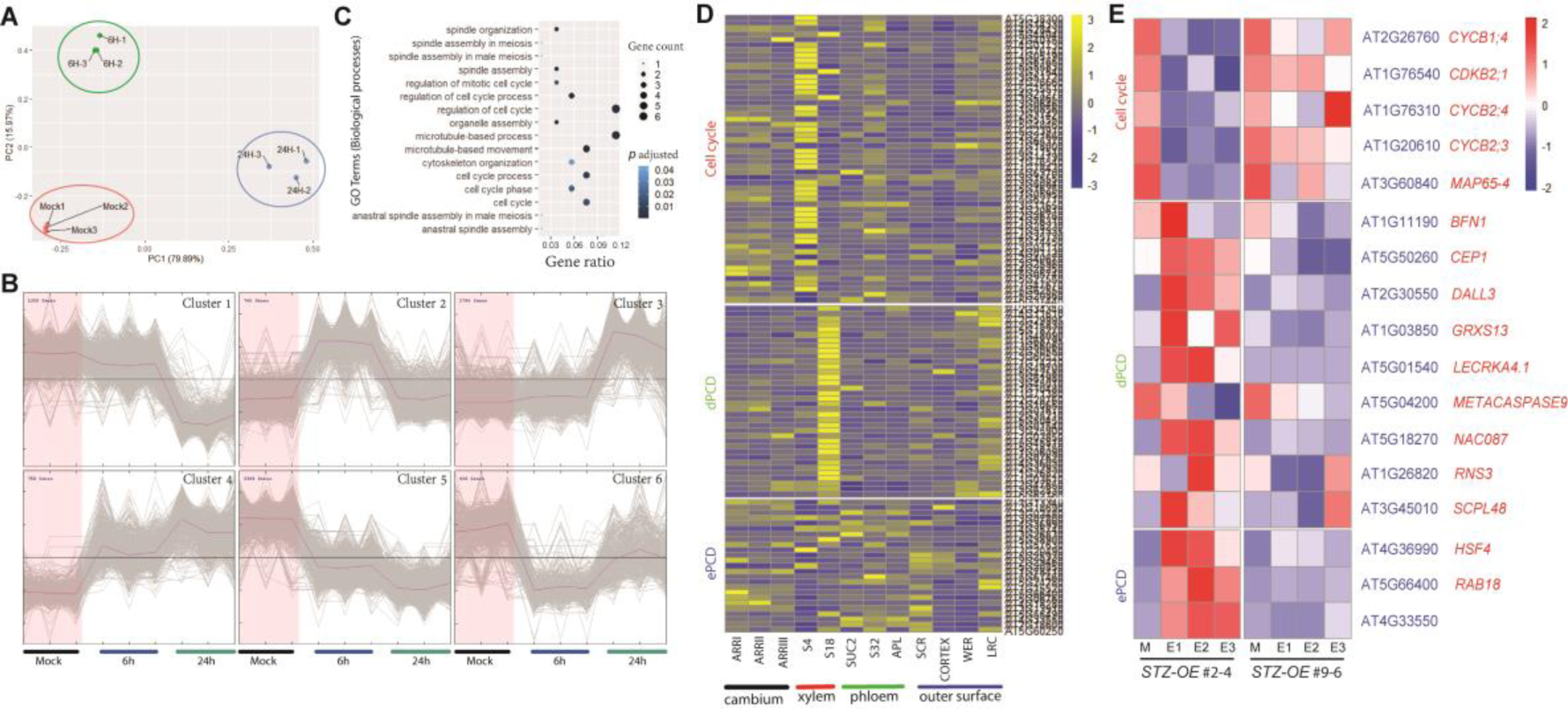
*STZ* transcriptionally regulates a group of cell cycle, developmentally induced programmed cell death (dPCD), and environmentally induced PCD (ePCD) genes downstream of stress signaling. **(A)** A principle component analysis (PCA) of the nine *STZ-OE* RNA-seq samples based on the top 5000 most variable genes. We used RNA samples from 2 different time points (6H and 24H) following *STZ* induction and one set of mock samples. The sample information and raw data are presented in table S12 and 13. **(B)** The K-means clustering of 8,435 DEGs was grouped into six different clusters based on the expression dynamics. The data are also presented in Table S15. **(C)** GO enrichment (biological processes) analysis (FDR < 0.05) of the common cell cycle target DEGs suppressed both by PEP1 treatment and STZ overexpression. The data are also presented in Table S16 and 17. **(D)** Expression of *STZ* downstream cell cycle, dPCD and ePCD genes as obtained in our RNA-seq in the selected cell types including cambium, xylem, phloem and ground tissue layers of Arabidopsis root (Table S18; (Zhang et al., 2019)). Data were gene-wise normalized and plotted using the Pheatmap package of R. The abbreviated information of the cell-type specific markers is in Table S26. Notably, the cell cycle genes are enriched in the developing xylem (S4), whereas the dPCD markers are accumulated in maturing xylem (S18). For the ePCD genes, no specific patterns of tissue-specific localization were observed. **(E)** The transcript levels of the selective cell cycle, dPCD, and ePCD genes in the *STZ-OE* #2-4 and #9-6 lines as validated by qRT-PCR. The expression fold-changes were determined in the root samples treated with 20µM estradiol for 24h compared to the mock (ethanol treatment for 24h). The expression values normalized by *UBQ10* are presented as ± SEM from three technical replicates in Table S20. For visualization, gene-wise normalized expression values from three technical replicates in qRT-PCR were used to generate the heatmap.

*STZ-OE* #2-5 line exhibited an early and ectopic xylem differentiation (Figure 3h and I; S11) similar to the seedlings treated with PEP1 (Dhar et al., 2021). Therefore, we inquired whether STZ induced developmentally-induced programmed cell death (dPCD) genes. We compared the DEGs from the STZ-induced clusters with a core set of 98 conserved dPCD marker genes that were identified previously in context to tracheary elements and lateral root cap differentiation (Lee et al., 2006; Olvera-Carrillo et al., 2015), and identified 36 shared dPCD genes (Figure S12e, Table S19). We also identified 26 environmentally induced PCD (ePCD) marker genes upregulated by STZ (Figure S12f, Table S19) out of 84 known ePCD genes upregulated by several shared biotic, genotoxic, and osmotic stresses (Olvera-Carrillo et al., 2015). The expression profile of dPCD (36/98) and ePCD (26/84) genes in our transcriptome data is available in Figure S12g and h. Furthermore, we observed that these STZ downstream dPCD genes are specifically enriched in the maturing xylem layer (S18). In contrast, the ePCD genes are sporadically expressed throughout various tissue layers of the root (Figure 4d, Table S18). These findings suggest that STZ controls the cell cycle and cell differentiation at various tissue layers of the root.

Given a significant difference in root growth and stem cell population maintenance between *STZ-OE* lines #2 and #9 (Figure 3, S7, S8, and S9), we hypothesized that STZ dosages independently modulate root growth and tissue differentiation. To test this hypothesis, we measured the expression of selected cell cycle, dPCD, and ePCD genes in the roots of *STZ-OE* #2-4 and #9-6 using qRT-PCR (Figure 4e). As per our expectation, the cell cycle genes exhibited more that ∼50% suppression at 24h post-estradiol treatment in the *STZ-OE* #2-4 roots as compared with the mock, whereas *STZ-OE* #9-6 maintained a transcription level similar to ∼80% of the mock (Figure 4e; Table S20). Among the dPCD genes, we monitored the presence of several well-characterized dPCD regulators including a *NAC087*, a *SERINE CARBOXYPEPTIDASE-LIKE 48* (*SCPL48*), a *GLUTAREDOXIN13* (*GRXS13*), a ribonuclease *RNS3* (Figure S12g, Table S19). In Arabidopsis, NAC46 and NAC87 was shown to be highly expressed in xylem tracheary element to mediate PCD via the dPCD executers, nuclease BFN1 and ribonuclease RNS3 (Olvera-Carrillo et al., 2015; Fujimoto et al., 2018; Huysmans et al., 2018). Therefore, we measured the transcription levels of total nine dPCD related genes, *NAC087*, *SCPL48*, *GRXS13*, *RNS3*, *BFN1*, *DAD1-LIKE LIPASE 3* (*DALL3*), *LECTIN RECEPTOR KINASE A4.1* (*LECRKA4.1*), *METACASPASE9* (*MC9*), and *CYSTEINE ENDOPEPTIDASE* (*CEP1*) in *STZ-OE* #2-4 and *STZ-OE* #9-6 (Figure 4e; Table S20). In general, the results indicated a higher induction of dPCD genes in *STZ-OE* #2-4 roots than *STZ-OE* #9-6 (Figure 4e; Table S20). By contrast, three ePCD marker genes, *HEAT SHOCK FACTOR* (*HSF4*), *ARABIDOPSIS THALIANA DROUGHT-INDUCED 8* (*RAB18*), and a bifunctional inhibitor/lipid-transfer protein (*AT4G33550*) were induced at variable levels by *STZ* (Figure 4e; Table S20). Taken together, these observations suggest that STZ, being activated by stress signals, influences cell cycle and differentiation gene cascades at the root apical meristem in a dosage-dependent manner.

### STZ dosage controls the magnitude of stress response and organ growth

The transcription of *STZ* started upon PEP1 perception, and the dosage gradually increased at 24h post-PEP1 treatment (Figure 2b). Using our *STZ-OE* lines, we artificially intended to mimic different dosage of *STZ* that cumulates following perception of stress signals. The seedlings of *STZ-OE* #2-5 and *STZ-OE* #6-3, showing very to moderately high dosage of *STZ*, failed to recover the root and shoot growth once the overexpression of *STZ* was ceased (Figure S9). However, *STZ-OE* #9-6 seedlings exhibited growth recovery to a level similar to WT (Figure S9). This observation hinted that plants might adjust STZ dosages to determine the degree of stress response and growth inhibition along the stress perception pathway.

In the PEP1-treated time-course transcriptome data, 39 well-known stress responsive TFs formed a network with *STZ* in the STRING database (Table S21; with STRING cut-off 0.4). Among these TFs, 19 TFs were DEGs in the *STZ-OE* data (Figure S12i, Table S22). Intriguingly, we observed these stress regulators are highly enriched in the cambium (ARRI), developing cambium (ARRII), companion cells (SUC2), phloem (APL), and cortex layer (CORTEX) in the root (Figure 5a).

**Figure 5.**
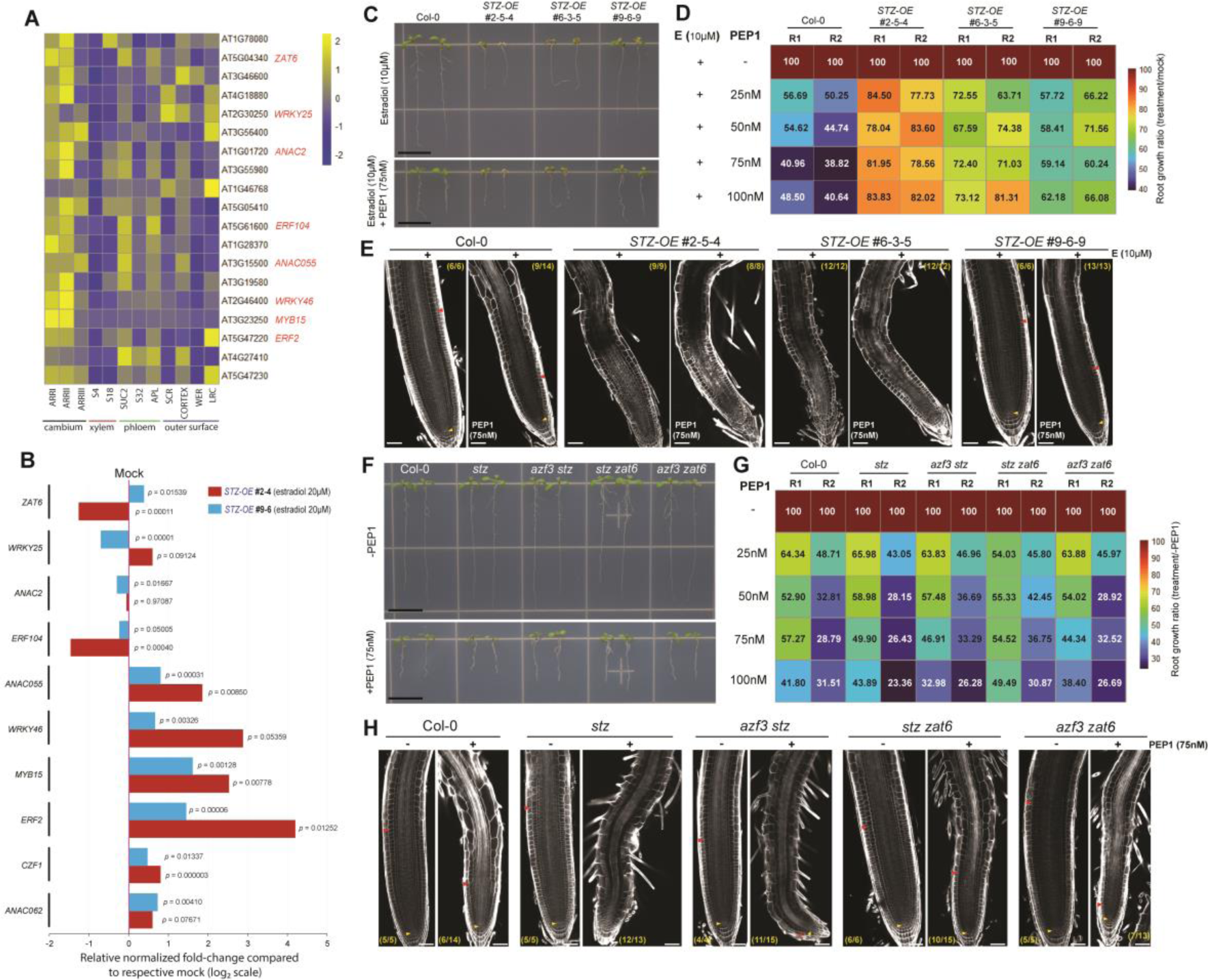
The magnitude of stress-responsive networks is differentially regulated by STZ at the root meristem. **(A)** The STZ-regulated stress genes are significantly enriched in the cambium and (ARRI) developing cambium (ARRII) tissue layer of the root (see also Table S18). The heatmap represents row-normalized (gene-wise) expression data. **(B)** The transcript levels of selective stress regulators in the *STZ-OE* #2-4 and #9-6 lines as validated by qRT-PCR (see also Table S20). *UBQ10* was used as an internal control, and the normalized fold-change was determined in the root samples treated with 20µM estradiol for 24h as compared with mock (24h ethanol treated). The bar graph represents the log_2_ |normalized fold-change| from three technical replicates. **(C)** The representative root growth images of 7 DAT old Col-0 and *STZ-OE* seedlings grown in the presence of 10 M estradiol and 75nM PEP1 for 4 days. The *STZ-OE* seedlings used in this experiment are the T4 generation from *STZ-OE* #2-5 (2-5-4), #6-3 (6-3-5), #9-6 (9-6-9). Scale bars = 10 mm. **(D)** Heatmap shows a relative root growth ratio of Col-0 and three *STZ-OE* lines compared to respective mock. Seedlings were grown on ½ MS media for 3 DAT, transferred to respective mock or treatment plates, and grown for 4 more days. ½ MS plates with only estradiol (10 M) were considered as mock, whereas the plates containing estradiol (10 M) and various dosages of PEP1 media (25, 50, 75, and 100 nM) were used as treatment. The quantitative growth data is also presented in Table S23. The numbers in each color blocks represents the percentage of root length in the respective treatment compared to mock. *STZ-OE* lines exhibited better root growth on PEP1 compared to Col-0. **(E)** The meristem growth of the Col-0 and *STZ-OE* lines under the condition mentioned in C. The yellow and red arrows indicate QC and meristem/transition zone boundaries, respectively. Numbers in parenthesis of each panel indicate samples with similar results of the total independent root samples analyzed. Scale bars = 30 m. **(F)** Representative root growth images of 7 DAT old Col-0 and *stz-*related mutant seedlings grown in the presence of 75 nM PEP1 for 4 days. Scale bars = 10 mm. **(G)** Heatmap represents a relative root growth ratio of Col-0 and *stz-*related mutants compared to the respective ½ MS grown roots. Seedlings were initially grown on ½ MS up to 3 DAT and transferred to respective -PEP1 or treatment plates (25, 50, 75 and 100 nM of PEP1) and grew for 4 more days. The quantitative growth data is also presented in Table S23. The numbers in each color blocks represents the percentage of root length in the respective treatment compared to root samples grown without PEP1. **(H)** The meristem growth of the Col-0 and *stz-*related mutants under the condition mentioned in **F**. The yellow and red arrows indicate QC and meristem/transition zone boundaries, respectively. Numbers in parenthesis of each panel indicate samples with similar results of the total independent root samples analyzed. Scale bars = 30 m.

Among 19 stress response TFs that are downstream of both PEP1 and *STZ-OE* #2-4, the expression of 8 well-known stress regulators, as well as two direct targets of STZ (O’Malley et al., 2016), was monitored in *STZ-OE* #2-4 and #9-6 roots using qRT-PCR (Figure 5b; Table S20). *WRKY46*, *MYB15*, *ANAC055*, and an *ETHYLENE RESPONSE FACTOR-2* (*ERF2*) were highly upregulated (> 4-fold compared to mock) in *STZ-OE* #2-4 roots (Figure 5b; Table S20). Though *STZ-OE* #9-6 root did not show root growth suppression upon STZ induction (Figure 3c), it displayed a significant increase (1.5∼3-fold compared to mock) of RNA levels for these key stress regulators. *CZF1* and *ANAC062*, direct targets of STZ (O’Malley et al., 2016), were also induced (Figure 5b; Table S20). Furthermore, the *ZAT6*, a close STZ homolog, was differentially regulated by STZ dosages, i.e. suppressed in #2-4 by activated in #9-6 (Figure 5b; Table S20). These observations suggested that STZ manipulates the magnitude of stress resistance at the root, depending on the *STZ* level. The basal level for inducing stress resistance seems lower than the level required for growth suppression since STZ-OE #9-6 activates stress resistance genes without suppressing the growth. Depending on the stress level, plants might set the STZ dosage level to decide the growth and defense tradeoff.

### Engineering STZ dosages for making stress-resilient plants

STZ differentially controls stress signals and growth in a dosage-dependent manner. Therefore, we investigated whether STZ doses differentially affect root growth under the influence of danger signals (Figure 5c - e). We monitored the root growth in lines #2-5-4, #6-3-5, and #9-6-9, which are T4 progenies of #2-5, #6-3, and #9-6, under the influence of mock (10 µM of estradiol) and various dosages of PEP1 (25nM, 50nM, 75nM, and 100 nM with 10 µM of estradiol). As expected, the root growth of *STZ-OE* #2-5-4 and #6-3-5 exhibited suppression of root growth, likely due to a higher dosage of *STZ*, whereas *STZ-OE* #9-6-9 showed root longer than lines #2-5-4 and #6-3-5 when grown on mock (Figure5c, Table S23). However, when we added PEP1, we observed a higher resistance in the suppression of root growth in *STZ-OE* #2-5-4 and #6-3-5 lines as they maintained a root growth around ∼77-84% (in #2-5-4) and ∼63-72% (in #6-3-5) compared to the mock (Figure 5c and d, Table S23). Furthermore, *STZ-OE* #9-6-9, which contains a higher dosage of *STZ* than Col-0, displayed resistance in root growth suppression (∼59-66% of the mock) as compared to Col-0 (∼38-48% of the mock), especially when the higher dosage of PEP1 (75nM and 100nM) was applied (Figure 5d, Table S23). When we profiled the exhaustion of meristematic cells in the above *STZ-OE* lines at mock and PEP1 treatment, #9-6-9 sustained proliferative cell activity better than Col-0, whereas a higher dosage of *STZ* together with PEP1 perception caused a complete exhaustion of meristem which leads to the loss of re-growth potential in the seedlings once stress had been removed (Figure 5e, S9, S15). These observations suggested that a higher dosage of *STZ* contributed to a higher resistance at the cost of growth. By contrast, a mild induction of *STZ* over the basal level in *STZ-OE* #9-6-9 provides better resistance in growth suppression than wild-type without compromising the re-growth potential.

To further characterize the variable role of STZ under stress, we generated mutant combinations of *STZ* homologs using the multiplex CRISPR/Cas9 assembly system combined with fluorescent seed selection (Ursache et al., 2021); (Figure S13). We introduced Cas9-gRNA for *AZF3* and *ZAT6* into the *stz* T-DNA insertion mutant (SALK_054092) to generate *azf3 stz* and *stz zat6* double mutants (Figure S14a and b; methods). *azf3 zat6* double mutant was identified in the screen for transgenic Col-0 introduced with the gRNAs targeting *AZF3* and *ZAT6* (Figure S14c; methods). To visualize the root growth sensitivity under various dosage of PEP1, we grew the seedlings of Col-0, *stz*, *azf3 stz*, *stz zat6*, and *azf3 zat6* on ½ MS plates for 3 days before they were transferred to either mock (-PEP1) or PEP1 containing media and further grown for 4 more days. With the increasing PEP1 concentration, a gradual suppression of root growth was observed in Col-0 (Figure 5f, Table S23). Given that there are differences between each biological replicates in terms of root growth suppression ratio, we tried to find the consensus observations between them. The *stz*, which lacks STZ-activated stress regulatory network, displayed a hyper sensitive in root growth suppression compared to Col-0 at higher dosage of PEP1 (75nM and 100nM). A differential root growth ratio was observed in *azf3 stz* and *azf3 zat6* double mutants compared to respective mock (Figure 5g; Table S23). Intriguingly, *stz zat6* showed a decline in the root growth ratio (∼54% (replicate 1) and ∼46% (replicate 2) as compared to mock) when exposed to a mild dosage of PEP1 (25nM), whereas in 50, 75 and 100nM of PEP1 treatment, we observed a better root growth to PEP1 compared to Col-0 (Figure 5g; Table S23). In addition, we found that the meristem of the *stz zat6* was sustained as compared with Col-0 and *stz* while most of the Col-0, and *stz* seedlings displayed exhaustion of the meristem following PEP1 exposure (Figure 5h; S15). Taken together, these observations suggested that *STZ* and *STZ*-like (*STZL*) genes redundantly regulate the stress responses and reprogram the growth potential of the meristem. However, further studies are essential to identify the complex interplay between these TFs that attributes to engineer stress resilient plants.

## Discussion

In nature, roots are under constant exposure to complex environmental stresses. These stresses turn on the endogenous danger signals to act for the growth and defense tradeoff (Huot et al., 2014; Figueroa-Macias et al., 2021). PEP1, a danger signal activated (Bartels and Boller, 2015) by various biotic/abiotic stresses, triggers defense responses while suppressing apical root and meristem growth and inducing root hair and vascular tissue differentiation (Poncini et al., 2017; Jing et al., 2019; Dhar et al., 2021). However, it remains elusive how and when PEP1 transcriptionally manipulates root developmental pathways and activates defense response (Bartels and Boller, 2015). In this study, we aimed to understand how the early point of stress perception affected the developmental outcome by using a combination of approaches, including genome-wide expression analysis, gene network-based functional inferences, various gene perturbations, and detailed phenotype analyses.

The root meristem is the central place governing growth; thus, it is likely that PEP1 manipulates developmental resources in the meristem to cope with exogenous danger signals. Indeed, our high-resolution meristem-specific RNA-seq. dataset revealed rapid induction of a plethora of stress-responsive TFs in the meristem as early as 1-hour post-PEP1 treatment. We assumed some of those TFs are responsible for the strong suppression of cell cycle genes, which maintain cell divisions and, thereby, the stem cell population in the meristem (Figures 1 and 2). Because rich information about networks amongst stress-responsive TFs and cell cycle regulators is available, we searched for the connection between the two categories downstream of PEP1 signaling using the STRING database. In this network analysis, *STZ* was predicted as one of the most highly connected TFs in the stress response network and connected to the cell cycle networks.

STZ, AZF3, and ZAT6, the most closely related members of the C2H2 zinc finger TF family, are all induced by PEP1. Expression dynamics of STZ, AZF3, and ZAT6 in response to PEP1 treatment further support their potential role in the regulation of the cell cycle network, inferred from our STRING database analysis. Their expression is strongly induced in the root apical meristem with PEP1, while no expression is found without PEP1. STZ and ZAT6 proteins are distributed at the root stem cell niche, indicating that PEP1-mediated reprogramming of root development might happen through STZ and related C2H2 zinc finger TFs. Indeed, expressing a high dosage of STZ in the root meristem using an estradiol-inducible transactivation system (*STZ-OE* line #2) resulted in (i) the exhaustion of stem cell niche, (ii) an early and ectopic xylem vessel differentiation, and (iii) aberrant divisions in the QC. These phenotypes are reminiscent of those found in the seedlings treated with PEP1 (Jing et al., 2019; Dhar et al., 2021; Okada et al., 2021). The *STZ-OE* transgenic line #2 phenotypes are further supported in the RNA-seq analyses. STZ-suppressed clusters were enriched with cell cycle genes, while the STZ-activated clusters were with dPCD-/ePCD-associated regulators and executors (Olvera-Carrillo et al., 2015) (Figure 4). Most dPCD-associated genes were found to be enriched in the maturing xylem cell, implying their role in the differentiation of xylem vessels.

Fine-tuning the growth and defense tradeoff is critical for plant sustainability in constantly changing environments (Karasov et al., 2017), particularly for the root system. Once the stress-responsive network is activated, the developmental resources are taken away. However, this study found that STZ regulates stress response and growth-related genes at different levels (Figure 5). Amongst estradiol-inducible *STZ-OE* lines, line #9 with the relatively low *STZ* induction level showed root and meristem growth comparable to the WT even after *STZ* induction; however, it exhibited activation of ANA055, MYB15, ERF2, ZAT6, CZF1, and *ANAC062* among the key stress regulators (Figure 5). The phenotype of STZ-OE line #9 contrasts with *STZ-OE* line #2, which exhibited a high dosage of stress gene induction and strong suppression of root growth (Figure 5). Consistent with the induction of key stress regulators, the roots of line #9 (#9-6-9; T4 generation) exposed to PEP1 grew longer than Col-0 without compromising meristem exhaustion (Figure 5). By contrast, stronger root growth suppression was found in *stz* mutant under PEP1 than in the WT. These findings collectively suggest the possibility of designing plants with enhanced stress resistance without compromising growth by modulating STZ levels.

In this study we revealed a transcriptional landscape of the root meristem upon perception of PEP1. We also demonstrated that *STZ* and its homologs are a potential nexus between danger signal perceptions and molecular reprogramming of the meristem. Our findings suggest that PEP1-mediated cell division suppression, cell differentiation induction, and activation of stress regulators in the root meristem depend on the dosage of *STZ*, which varies based on the exposure or magnitude of stress signals (Figure 6).

**Figure 6.**
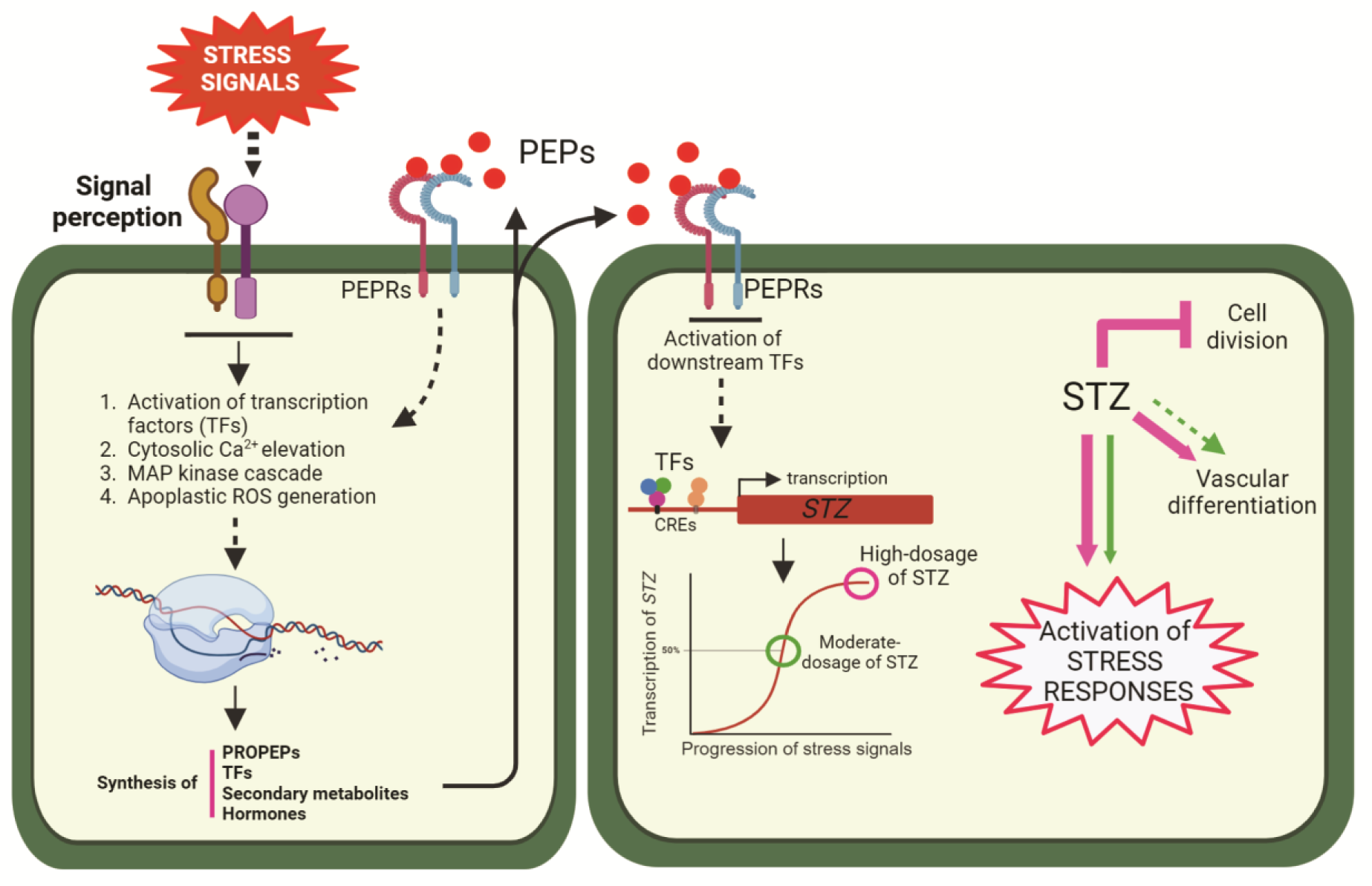
A working model (created with BioRender.com) showing the perception/progression of stress signals (Yamaguchi and Huffaker, 2011), which are directed to developmental reprogramming. This study used synthetic PEP1 as a representative candidate for stress-induced DAMPs. PEPs, secreted in response to exogenous stress signals, activate a group of TFs, including C2H2 zinc fingers, which serve as a potential nexus where danger signals converge and modulate the developmental program. The magnitude of stress signals controls transcript levels of STZ, which differentially regulates stress responses and growth in the root meristem. In our observation, a high dosage of STZ strongly activates stress resistance while compromising root meristem activity and growth. By contrast, a moderate dosage of STZ activates a stress resistance network without compromising root growth, which might be beneficial for sustaining plant yield under a moderately harsh environment.

## Materials and methods

### Plant materials and growth conditions

The Arabidopsis plants used in this study were the Columbia (Col-0) ecotype. Col-0 plants were used as the wild-type (WT) control in this study. The mutants *stz* (SALK_054092) (Hoang et al., 2020), *pepr1 2* (Nakaminami et al., 2018), and transgenic lines *ProWOX5:erGFP* (Sebastian et al., 2015); *ProTMO5::erGFP* (Lee et al., 2006), and PlaCCI (plant cell cycle indicator) (Desvoyes et al., 2020) were reported previously. All seeds were surface-sterilized, vernalized and grown on ½ MS media supplemented with 1% sucrose under a 16-h-light/8-h-dark cycle with 100 μmol m−^2^ s−^1^ light intensity at 22°-23°C in a plant growth chamber.

### Root and meristem growth suppression assay

The synthetic PEP1 (ATKVKAKQRGKEKVSSGRPGQHN) was obtained from Peptron (Korea) (http://www.peptron.com) (Dhar et al., 2021). The peptide was dissolved in distilled water to make stocks of 10mM each and stored in a -20°C freezer until use.

To analyze the root growth in *STZ-OE* lines, we surface sterilized seeds and plated the respective lines of *STZ-OE* into ½ MS plates supplemented with 10 M estradiol (Sigma-Aldrich, USA). The plates were cold treated and grown at 22°C plant growth chamber under long-day condition up to 5 days after transfer to the growth chamber (DAT). For the root growth suppression phenotype in *STZ-OE* and *stz* related mutants, seedlings grown on solid ½ MS plates were transferred at 3 DAT condition to fresh ½ MS plates supplemented with various concentrations of PEP1 and grew them for four more days as mentioned in the respective figures. The plates were photographed using a Nikon digital camera, and root length was measured using ImageJ (https://imagej.net/ij/).

To visualize meristem growth, we harvested and fixed the seedlings into 4% paraformaldehyde (PFA) overnight. The seedlings were washed using 1X phosphate-buffered saline (PBS) (137 mM NaCl, 2.7 mM KCl, 10 mM Na_2_HPO_4_, 1.8 mM KH_2_PO_4_; pH 7.4) and treated with ClearSee (Kurihara et al., 2015) for 3 to 5 days. Seedlings were stained with 0.1% (v/v) calcofluor white 2MR (Sigma-Aldrich, USA) for at least 1 hour and visualized using a confocal microscope (Leica TCS SP8) with preset excitation/emission wavelength of calcofluor white 2MR (350 nm/420 nm).

### Tissue harvest, RNA extraction and sequencing

For PEP1 time-course transcriptome data, 4-5 DAT seedlings of Col-0 were treated with (PEP1) or without (mock) 1 M PEP1 for 24h, 12h, 6h and 3h in such a way that at the sample harvesting time the age of the seedlings was 5 DAT. The root meristems were dissected from each set of seedlings, and the tissues were initially fixed overnight using an RNA*later* (Sigma-Aldrich, USA) solution. The total RNA was extracted using a RNeasy® Plant Micro Kit (Qiagen, Germany) with an on-column DNase digestion step to remove genomic DNA contamination.

For preparing RNA samples for time-course *STZ-OE* RNA-seq. data, 4-5 DAT seedlings of *STZ-OE* #2-4 were treated with equal volume of ethanol (mock) or 20 M estradiol for 24h and 6h, respectively. Approximately the bottom half of the roots were dissected from the 5 DAT seedlings, and total RNA was isolated using RNeasy® Plant Mini Kit (Qiagen, Germany) with an on-column DNase digestion step following manufacturer instructions.

The RNA quality, integrity, and quantity were measured using a Nanovue spectrophotometer (GE Healthcare, USA) and by a 2100 Agilent Bioanalyser with a plant RNA NanoChip assay (Agilent Technologies, USA) (Table S24). The library was prepared with the TruSeq Stranded mRNA Library Prep Kit (Illumina, USA), having an average insert size of > 300 bp. The NGS library for each sample was sequenced using an Illumina NovaSeq6000 instrument to obtain 100-bp paired-end reds at Macrogen Inc. (Seoul, Korea). The sequencing files have been deposited to the Gene Expression Omnibus (http://www.ncbi.nlm.nih.gov/geo/) with the GEO accession numbers GSE252414 (PEP1 RNA-seq) and GSE252415 (*STZ-OE* RNA-seq).

### Data analysis for RNA-seq

The read quality of the obtained FASTQ files were assessed by FastQC (https://www.bioinformatics.babraham.ac.uk/projects/fastqc/). Trimming of adapter sequences and low-quality reads was done using CLC genomics workbench v 22.0.1 (Qiagen, Germany) with a parameter-quality limit of 0.05. To estimate transcript abundance, high-quality trimmed reads were mapped to the Arabidopsis genome (TAIR10) with the parameters-mismatch cost: 2, insertion cost: 3, deletion cost: 3, length fraction: 0.8, similarity fraction: 0.8, and the expression level was normalized as reads per kilobase per million mapped reads (RPKM) at CLC genomics workbench. The DEGs (FDR < 0.01 and |fold change| ≥ 1.5) were determined by comparing the PEP1 or estradiol-treated samples with their respective mock.

### PCA, Clustering, GO, and KEGG pathway analysis of the DEGs

For PCA analysis, the top five thousand variably expressed genes were used among the DEGs. R package “ggfortify” and “ggrepel” were used to visualize the PCA data. The DEGs were combined and K-means clustered using the MeV software v4.9.0 (Saeed et al., 2003). GO term enrichment analyses were done via a hypergeometric distribution test on a comparison against the Arabidopsis reference genes as the background. GO and GOSlim categories were assessed with the BINGO package (Maere et al., 2005) in Cytoscape (www.cytoscape.org), keeping the P-value of ≤ 0.05. The data presentation was done in RStudio using “ggplot2” package (Villanueva and Chen, 2019).

### Gene expression quantification by qRT-PCR

Around 2 µg of purified RNA was used as a template for cDNA biosynthesis using SuperScript™ III Reverse Transcriptase (Invitrogen, USA) in a 20 µl reaction. The synthesized cDNA was diluted 5-fold by adding 80 µl of autoclaved distilled water, and 1 µl of cDNA was used as a template for qRT-PCR using iQ^TM^ SYBR^®^ Green Supermix (BIO-RAD, USA) on a CFX96^TM^ Real-Time PCR machine (BIO-RAD, USA) by following manufacturer’s instruction as previously reported (Kim et al., 2020; Dhar et al., 2021). The sequence information of gene-specific primers is in Table S25.

### Molecular cloning and generation of transgenic plants

To generate transcriptional reporter lines of *STZ*, *AZF3,* and *ZAT6*, MultiSite Gateway® system (Invitrogen, USA) was used as previously described (Lee et al., 2006; Kim et al., 2020). The promoter regions of *STZ* (Sakamoto et al., 2004), *AZF3* (Sakamoto et al., 2004), and *ZAT6* (Shi et al., 2014) were amplified from Col-0 genomic DNA by PCR and inserted into pDONR P4-P1r via BP recombination. The reporter component *eGFPGUS* (Furuta et al., 2014) was cloned into pDONR221. To obtain *pSTZ::eGFPGUS*, *pAZF3::eGFPGUS* and *pZAT6::eGFPGUS*, final recombination was done into the binary vector dpGreenBarT using Multisite Gateway LR recombination. To create the translational fusion lines of *STZ* (*pSTZ::STZ-GFP*) and *ZAT6* (*pZAT6::ZAT6-GFP*), the coding sequences were amplified using Col-0 cDNA and inserted into the binary vector dpGreenBarT with the previously prepared cassettes of respective promoters at pDONR P4-P1r and GFP at pDONR P2R-P3.

To generate the conditional overexpression lines of *STZ* (*STZ-OE*), the *STZ* coding region was cloned into pENTR™ /D-TOPO™ vector (Thermo Fisher Scientific, USA). The pDONR P4-P1r vector containing *p35S::XVE>>pLexA* was obtained from Dr. Ari Pekka Mähönen (Siligato et al., 2016). The *p35S::XVE>>pLexA::STZ* was constructed into dpGreenBarT by LR recombination. All clones in the binary vector were introduced to Agrobacterium GV3101 with pSOUP for Arabidopsis transformation by floral dipping (Clough and Bent, 1998). The sequence information of the primers used for cloning are in Table S25.

### Generation of CRISPR-based mutant lines

We used the CRISPR-Cas9 technology to generate single and double mutant combinations (Ursache et al., 2021). The guide RNA (gRNA) was designed using a web-based program CRISPR-PLNT (www.omap.org/crispr) (Xie et al., 2014). Since the coding sequences are small (ZAT6= 717 bp, AZF3= 582 bp), for each gene, we designed two gRNAs (Table S25). The first and second gRNAs were cloned to the pRU41 and pRU42 vectors, respectively, following the method previously described (Ursache et al., 2021). Golden gate assembly of the gRNAs was performed to generate a single cassette containing both guide RNAs into the vector pSF463 (Ursache et al., 2021). Finally, the gRNA cassettes containing two gRNAs were switched to Gateway-compatible pRU53 (*AZF3* gRNA) and pRU54 (*ZAT6* gRNA) vectors using LR recombination. A schematic diagram of the cloning is in Figure S13.

To generate the *stz azf3* and *stz zat6* double mutants, the *AZF3* (pRU53 with FastRed) and *ZAT6* (pRU53 with FastGreen) gRNA-Cas9 transgenes were introduced into *stz* mutant through floral dipping. T1 generation was screened under INCell 2000 (GE health care) analyzer based on fluorescence. To generate the *azf3 zat6* double mutant, the gRNAs for *AZF3* (pRU53 with FastRed) and *ZAT6* (pRU54 with FastGreen) were introduced to wild-type Col-0 plants. The primary selection at the T1 generation was done based on the seeds having both FastRed and FastGreen signals. The seedlings used for the experiment are T3 generation containing Cas9 in the genome. To confirm the nucleotide changes, the target regions were amplified from genomic DNA of the transgenic plants. The sequencing results are available in Figure S14, and the primers used for this experiment are in Table S25.

### Imaging of transcriptional and translational marker lines

To identify the expression patterns of transcriptional and translational reporter lines, 4 DAT seedlings were treated with 1 M PEP1 for 24h before they were imaged using confocal miscrscopes (Leica TCS SP8 and Zeiss LSM700). To visualize the GFP protein, seedlings were stained in 1 µM propidium iodide (PI, Life Technologies) solution for 1 min and observed with the following excitation and detection windows: GFP 488 nm/500-530 nm; PI 555 nm/591-635 nm. For Z-scan imaging, the same position of the root and the same laser scanning area were selected. The GUS staining was performed following standard methods (Jefferson et al., 1987), and seedlings were imaged using a digital microscope (Nikon, Japan) with DIC optics.

### Histological analysis

To analyze the stele internal tissue organization, 5 DAT seedlings grown on half-strength MS media supplemented with 10 M estradiol were used. Five to six seedlings were arranged within a 4% low-melting SeaPlaque®, Agarose (Lonza) blocks. The solidified agarose was dissected into 120∼130 m slices containing root sections using a vibratome (Leica VT1000S). For WT (MS), WT (estradiol) and *STZ-OE* #2-5 (MS) samples, sections were made at the 4-6 mm basal region from the root tip and imaged using a confocal microscope (Zeiss LSM700) as previously reported (Kim et al., 2020; Seo and Lee, 2021). Due to the tiny root in *STZ-OE* #2-5 (estradiol), we performed plastic sectioning to determine the internal tissue organization. The seedlings were harvested and fixed overnight in 4% PFA at room temperature. The samples were then washed in 1X PBS, dehydrated in an ethanol series (30%, 50%, 70%, 90%, and 100% (v/v)), and eventually, the plastic blocks were prepared with Technovit 8100 kit by following instructions of the manufacturer. As described previously, 5 m width plastic sections were made within the 1 mm basal region from the root tip using a Leica RM2255 microtome (Kim et al., 2020; Dhar et al., 2021). The sections were imaged using a Nikon Eclipse N*i*-U microscope with DIC optics and presented in Figure S11.

### Quantification and statistical analysis

All statistical analyses, unless otherwise stated, were performed using Microsoft Excel. Plots and graphs were generated using the R program and RStudio software (https://www.rstudio.com) using ggplot2, tidyverse and pheatmap packages, and Excel. Venn diagrams were drawn using online tools (http://bioinformatics.psb.ugent.be/webtools/Venn). All analyses in CLC-Genomics Workbench (Qiagen, Germany) and the Linux environment were conducted on a local server hosted by the Plant Systems Genetics lab at Seoul National University, Korea.

### Accession numbers

The RNA-seq. data reported in this work have been submitted to the NCBI GEO database (http://www.ncbi.nlm.nih.gov/geo) with the accession numbers GSE252414 (PEP1 RNA-seq.) and GSE252415 (*STZ-OE* RNA-seq.). The sequence information of the genes used in this study can be found in the Arabidopsis Genome Initiative under the following accession numbers: *STZ* (AT1G27730), *AZF3* (AT5G43170), *ZAT6* (AT5G04340), *PEPR1* (AT1G73080), *PEPR2* (AT1G17750), *CYCB1;4* (AT2G26760), *CDKB2;1* (AT1G76540), *CYCB2;4* (AT1G76310), *CYCB2;3* (AT1G20610), *MAP65-4* (AT3G60840), *BFN1* (AT1G11190), *CEP1* (AT5G50260), *DALL3* (AT2G30550), *GRXS13* (AT1G03850), *LECRKA4.1* (AT5G01540), *MC9* (AT5G04200), *NAC87* (AT5G18270), *RNS3* (AT1G26820), *SCPL48* (AT3G45010), *HSF4* (AT4G36990), *RAB18* (AT5G66400), AT4G33550, *WRKY25* (AT2G30250), *ANAC2* (AT1G01720), *ANAC055* (AT3G15500), *WRKY46* (AT2G46400), *MYB15* (AT3G23250), *ERF2* (AT5G47220), *ERF104* (AT5G61600), *CZF1* (AT2G40140), *ANAC062* (AT3G49530), *WOX5* (AT3G11260), *TMO5* (AT3G25710), *GAPDH* (AT1G13440), *UBQ10* (AT4G05320).

## Data availability statement

All the data and materials that support the findings related to this study are included as a part of the article or Supplemental materials.

## Acknowledgements

We thank the members of the Lee lab for assisting in the experiments at various stages. We are grateful to Dr. Kousuke Hanada for generously providing *pepr1 2* seeds. This work was supported by the grants NRF-2021R1A2C3006061 and NRF-2018R1A5A1023599 to J.Y.L. from the National Research Foundation of Korea. S.D., J.P., and H.S. were supported by the Brain Korea 21 Plus Program. J.P. was supported by Dr. Hyung & Gertruda Choe Scholarship and Fellowship for Fundamental Academic Fields at Seoul National University.

## Author Contributions

Conceptualization, S.D. and J.Y.L.; methodology, S.D., S.K., and J.Y.L.; investigation, S.D., S.K., and J.Y.L.; writing – original draft, S.D.; writing – review & editing, S.D. and J.Y.L.; funding acquisition, J.Y.L.; resources, S.D., S.K., H.S., J.P., and J.Y.L.; supervision, J.Y.L.

